# Measuring metabolic rate in single flies during sleep and waking states

**DOI:** 10.1101/2021.11.10.468156

**Authors:** Elizabeth B. Brown, Jaco Klok, Alex C. Keene

## Abstract

*Drosophila melanogaster* is a leading genetic model for studying the neural regulation of sleep. Sleep is associated with changes in behavior and physiological state that are largely conserved across species. The investigation of sleep in flies has predominantly focused on behavioral readouts of sleep because physiological measurements, including changes in brain activity and metabolic rate are less accessible. We have previously used stop-flow indirect calorimetry to measure whole body metabolic rate in single flies and have shown that in flies, like mammals, metabolic rate is reduced during sleep. Here, we describe a modified version of this system that allows for efficient and highly sensitive acquisition of CO_2_ output from single flies. We also describe a modification that allows for simultaneous acquisition of CO_2_ and O_2_ levels, providing a respiratory quotient that quantifies how metabolic stores are utilized. Finally, we show that sleep-dependent changes in metabolic rate are diminished in aging flies, supporting the notion that sleep quality is reduced as flies age. Taken together, the use of indirect calorimetry provides a physiological measure of sleep with broad applications to genetic studies in flies.

## INTRODUCTION

Dysregulation of sleep is associated with weight gain, increased appetite, and numerous metabolism-related diseases including obesity, type II diabetes, heart disease, and metabolic syndrome (Aurora & Punjabi, 2013; Leproult & Van Cauter, 2010; Reutrakul & Van Cauter, 2014; Sharma & Kavuru, 2010). While insomnia is typically associated with reduced sleep duration, growing evidence suggests dysregulation of sleep quality has severe negative health consequences (Reutrakul & Van Cauter, 2018). In mammals, slow wave sleep is associated with a reduction in body temperature and glucose utilization, resulting in lower metabolic rate (Sharma & Kavuru, 2010). The metabolic changes associated with sleep are thought to be critical for the restorative effects of sleep on brain and endocrine function, however, little is known about the mechanisms underlying sleep-dependent changes in metabolic regulation (Zhang et al., 2002).

In recent decades, significant progress had been made identifying genes and neurons that regulate sleep (Artiushin & Sehgal, 2017; Chakravarti et al., 2017; Ly et al., 2018; Shafer & Keene, 2021). Forward genetic approaches in non-mammalian model systems including *C. elegans, D. melanogaster*, and *D. rerio* have provided critical insights into the conserved genetic and neural mechanisms governing sleep regulation (Hendricks et al., 2000; Prober et al., 2006; Raizen et al., 2008; Shaw et al., 2000; Yokogawa et al., 2007; Zhdanova et al., 2001). However, surprisingly little is known about sleep-dependent changes in metabolic rate. In mammals, temperature and O_2_ utilization provide measures of metabolic rate during waking and sleeping states and reveal elevated metabolic rate during periods of sleep deprivation and reduced metabolic rate during sleep (Caron & Stephenson, 2010; Katayose et al., 2009; Koban & Swinson, 2005; White et al., 1985). The combination of similar approaches in genetic model systems would provide new avenues to investigate interactions between sleep and metabolic rate.

Sleep in *Drosophila* is modulated by numerous environmental and life history traits including social context, feeding state, and age (Ganguly-Fitzgerald et al., 2006; Keene et al., 2010; Koh et al., 2006; Murphy et al., 2016). However, the effects of these conditions on sleep quality, and more specifically, sleep-dependent regulation of metabolic state, remain poorly understood. Towards this end, we previously developed a Sleep and Activity Metabolic Monitor (SAMM) system that simultaneously measures activity and metabolic rate using indirect calorimetry (Brown et al., 2019; Botero et al., 2021; Stahl et al., 2017; Stahl et al., 2018; Zandawala et al., 2018). This system uses stop-flow respirometry to measure CO_2_ output from individual flies during sleeping and waking states. While the SAMM system can detect circadian changes in metabolic rate as well as reductions in metabolic rate associated with prolonged sleep bouts, the sensitivity of the system remained a major impediment. Here, we report a modified system with increased sensitivity, allowing for the identification of changes in metabolic rate that result from genetic or environmental manipulations. We also apply the SAMM system to quantitatively assess the impact of aging on metabolic rate. We find that aging specifically interferes with sleep-dependent changes in metabolic rate, suggesting that aging in flies, like in humans, impairs sleep-associated changes in metabolism. Together, these findings support the notion that the SAMM system can be used to investigate novel integrators of sleep and metabolic state.

## METHODS

### *Drosophila* stocks and husbandry

The *w*^1118^ fly strain was used for all experiments (Bloomington Stock #5905; Ryder et al., 2004). Flies were grown and maintained on standard food media (Bloomington Recipe, Genesee Scientific). Flies were maintained in incubators (#DR-36VL, Percival Scientific) at 25°C on a 12:12 light/dark cycle, with humidity of 50%. Flies were isolated in groups at 1-2 days old, separated by sex at 2-3 days old, and then maintained on standard food media until testing. For aging experiments, flies were flipped every second day to fresh food. Unless otherwise noted, all flies were fed 1% agar (#BP1423, Fisher Scientific, Fisher Scientific) plus 5% sucrose (#S3, Fisher Scientific) during experimental trails.

### Measurement of CO_2_ output

The system includes 5 primary components (Figure 1). (1) A pump and scrubbing column, which removes CO_2_ coming into the system. (2) Two mass-flow control valves, which regulates air flow through the system. (3) A *Drosophila* locomotor activity monitor (DAM), which houses behavioral chambers and records activity of individual flies. (4) A multiplexer, that systematically directs accumulated air from an individual behavioral chamber to the (5) CO_2_ analyzer, which measures the CO_2_ produced. Each component is described in detail below:

**Figure 1.**
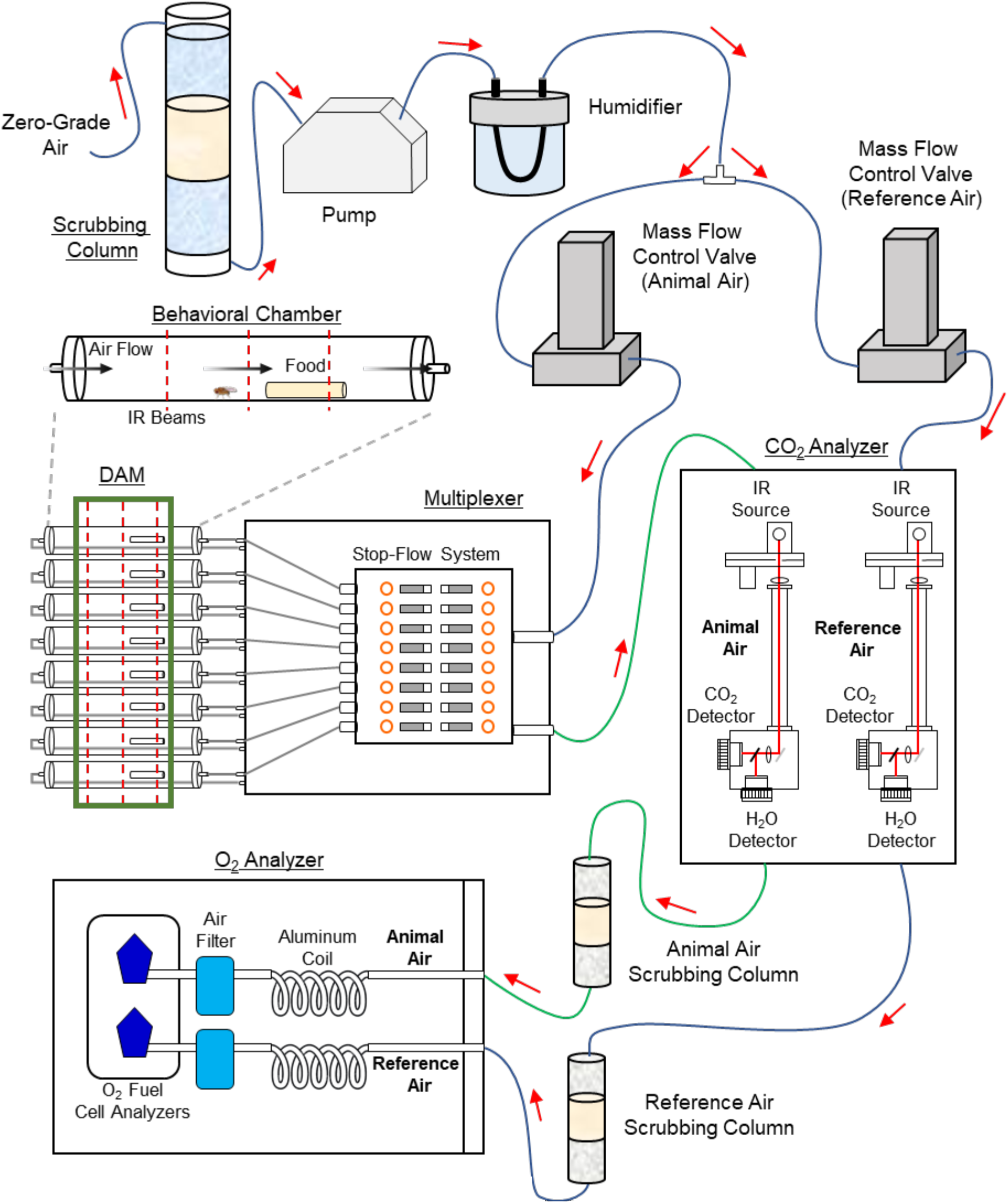
The Sleep, Activity, and Metabolic Monitor (SAMM) system can be used to measure sleep and CO_2_ output in single flies. Individual flies were fed 5% sucrose, during which sleep and CO_2_ output was measured over the course of 24 hours. To remove environmental CO_2_, ambient air is first passed through a scrubbing column. The resulting dehumidified, CO_2_-free air is then re-humidified and pumped to two mass-flow control valves to maintain constant air flow into the system. One mass flow control valve regulates clean, reference air flow directly to the CO_2_ analyzer. The other mass flow control valve regulates air flow to the multiplexer, which directs the sampling of CO_2_ that accumulates in each behavioral chamber over a specified time-period. The behavioral chambers are housed inside a *Drosophila* Activity Monitor (DAM), allowing for the simultaneous measurement of CO_2_ output and activity/sleep. The accumulated air from each behavior chamber is then sent to the CO_2_ analyzer, where the reference air is subtracted from the air collected from each behavioral chamber to determine CO_2_ output for each individual fly. Both the reference and animal air were then passed individually through scrubbing columns before entering the O_2_ analyzer, where the reference air is again subtracted from the air collected from each behavioral chamber to determine O_2_ consumed for each group of flies.

#### (1) Pump and scrubbing column

Nonpermeable Bev-A-line tubing (#56280, United States Plastic Corp.) was used throughout to connect each part of the system. This tubing was connected to a pump (SS-4 gas analyzer subsampler, Sable Systems International), pushing air into the respirometry system. To remove trace amounts of CO_2_ entering the system, zero-grade air, which contains less than 0.1 ppm total hydrocarbons, was passed through a scrubbing column (#26800, Drierite) containing an inner layer of ascarite (#81133-20-2, Acros Organics) and two outer layers of drierite (#7779-18-9, Drierite), each separated by a layer glass wool (#11-388, Fisher Scientific). To re-humidify, the air was then passed through water-permeable Nafion tubing (#TT-070, Perma Pure, LLC) immersed in a reservoir containing deionized H_2_O. The air was re-humidified to the dewpoint equivalent to the temperature chamber setting – 25°C.

#### (2) Mass-flow control valves

The re-humidified, CO_2_-free air was then split and directed to two mass flow control valves (Side-Trak 840 Series, Sierra Instruments, Inc.), both maintaining an experimental flow rate of 100 mL/min. One mass flow control valve regulates animal air flow into the multiplexer and DAM, while the other mass flow control valve regulates reference air flow directly into the CO_2_ analyzer’s reference channel.

#### (3) DAM

Behavior chambers were made of 70 mm × 20 mm custom-made glass tubes designed to fit inside a custom-made DAM (Trikinetics). The DAM contained three sets of infrared beams and was connected to a computer, where the number of beam-breaks by the flies were recorded every minute. To be able to assess baseline CO_2_ levels, one behavior chamber contained no flies. For trials with individual flies, 5 additional experimental behavior chambers were used. For trials with multiple flies, 7 additional behavior chambers were used.

#### (4) Multiplexer

The behavior chambers were connected to a stop-flow, push-through multiplexer air flow management system (RM8 Intelligent Multiplexer, Sable Systems International), thereby enabling the timed flow of air from one chamber to the next. While one chamber has active airflow all others are sealed off to accumulate fly respiratory gasses. For analysis of single flies, air was flushed from each of 6 behavior chambers for a duration 50 seconds. This provided a readout of accumulated CO_2_ output every 5 minutes. Sleep in *Drosophila* is typically defined by 5 min of behavioral quiescence (Shaw et al., 2000), thereby enabling sleep and activity metrics to be paired with CO_2_ output. For analysis of multiple flies, air was flushed from each of 8 behavioral chambers for a duration of 72 seconds, with the baseline being flushed twice, thereby providing a readout of accumulated CO_2_ output every 10 min. The baseline was measured twice so as to increase the sensitivity of O_2_ consumption measurements (see below), which was concurrently measured in groups of flies. These settings were applied using ExpeData PRO software (v1.9.27, Sable Systems International).

#### (5) CO_2_ analyzer

CO_2_ output was measured through indirect calorimetry using a CO_2_ analyzer (Li-7000, LI-COR). Clean, reference air was pumped directly into the CO_2_ analyzer independently from the animal air, which was pumped into the CO_2_ analyzer via the multiplexer.

All instruments were connected to a computer, where the temperature and humidity of the incubator as well as the flowrate, CO_2_ levels, and fly activity of the system were monitored. Flies were aspirated into behavior chambers ∼24 hrs prior to the start of each experiment to allow the flies to acclimate to the system. The respirometry system was calibrated every 3-5 trials. To set a baseline (CO_2_ = 0), the system was calibrated using pure Nitrogen. To calibrate CO_2_ levels, CO_2_ of a known concentration was used. Both N_2_ and CO_2_ were run though the system for a minimum of 1 hr.

### Measurement of O_2_ consumption

Once CO_2_ was measured in groups of 25 flies, air was passed from the CO_2_ analyzer to an O_2_ analyzer (Oxzilla II Oxygen Analyzer, Sable Systems International). Since water vapor dilutes oxygen content, the air was first scrubbed of all water vapor by passing through scrubbing columns. Two columns were used, one for the reference air and one for the animal air. Each scrubbing column composed the body of 10 mL syringe (#14955459, Fisher Scientific) containing an inner layer of ascarite and two outer layers of magnesium perchlorate (M54, Fisher Scientific), each separated by a layer glass wool. Rubber stoppers (#14-135E, Fisher Scientific) held the Bev-A-line tubing in place. These scrubbing columns were replaced twice over a 24 hr trial. The scrubbed air was then sent to the O_2_ analyzer for quantification.

### Analysis of CO_2_ output in single flies

CO_2_ output was analyzed using ExpeData PRO software (v1.9.27, Sable Systems International). For analyses of CO_2_, the absolute CO_2_ levels (ppm) extracted from each chamber was first transformed. To reduce noise, the data was smoothed using the Savitsky-Golay filter, with a window of 15. Next, lag correction of 8 seconds was applied, since there was a lag time of 8 seconds from when air flow exited the behavioral chamber to when it was measured by the CO_2_ analyzer. Last, to remove nonlinear drift, marker-pair baseline correction, with the Catmull-Rom Spline option was performed. To calculate VCO_2_, the absolute CO_2_ levels were converted to fractional content by dividing by 1,000,000, and then the fractional CO_2_ content was multiplied by the experimental rate of air flow to yield VCO_2_ in ml/min. This VCO_2_ data was then exported to Excel (Microsoft Windows), where the mean VCO_2_ values obtained from the baseline (empty) chamber was subtracted from each behavioral chamber for each 5 min time point. Next, the adjusted CO_2_ values were matched to their corresponding activity values (described in next section), thereby providing a direct comparison of CO_2_ output during wake and sleep. Here, we show VCO_2_ measurements in μL/hr, which was calculated by summing the 5 min time points into 1 hr bins.

### Analysis of sleep

Sleep and activity were measured using the *Drosophila* Locomotor Activity Monitor System, as previously described (Shaw et al., 2000). The DAM system measures activity by counting the number of infrared beam crossings for each individual fly. Since our DAM monitor had three infrared beams, overall activity was measured using a custom-generated python program that summed the raw activity from all three beams at each time point (Stahl et al., 2017) These activity data were then used to calculate bouts of immobility of 5 min or more using the *Drosophila* Sleep Counting Macro (Pfeiffenberger et al., 2010), from which sleep traits were then extracted.

### Analysis of CO_2_ output as a function of bout length

To investigate how CO_2_ output may change with sleep duration, we measured percent change in VCO_2_ over the duration of a single sleep bout. The change in VCO_2_ was calculated using the following equation: [(VCO_2_ @ 5 min) – (VCO_2_ @ 10 min)] / (VCO_2_ @ 5min) × 100. This was repeated for each 5 min bin of sleep, for the entire length of the sleep bout. Given that a single fly typically has multiple sleep bouts, the change in VCO_2_ for each 5 min bin of sleep was averaged across all sleep bouts over the course of the day/night. Additionally, we restricted our analysis to sleep bouts of up to 60 min, due to a decreased sample size of bouts of extended length.

### Analysis of CO_2_ output and O_2_ consumption in groups of flies

CO_2_ output was analyzed similarly as described above, with the exception that VCO_2_ values were obtained for each 10 min timepoint. For analyses of O_2_, both the reference air and the animal air extracted from each chamber were transformed. The data were first smoothed using the Savitsky-Golay filter with a window of 15 seconds. A lag correction of 30 seconds was then applied, since there was a lag time of 30 seconds from when air flow exited the behavioral chamber to when it was analyzed by the O_2_ analyzer. Last, a baseline correction was performed by correcting reference O_2_ to 20.95%, or a fractional value of 0.2095, which is the percent of O_2_ in dry air. VO_2_ was calculated from fractional contents using the following equation: (animal O_2_ – reference O_2_) × flow rate / (1 – reference air O_2_). The VO_2_ data was then exported to Excel, where the mean VO_2_ values obtained from the baseline (empty) chamber was subtracted from each behavioral chamber for each 10 min timepoint. Here, we show O_2_ measurements in μL/hr, which was calculated by summing the 5 min time points into 1 hr bins. For weight-adjusted measurements of CO_2_ output and O_2_ consumption, the VCO_2_ and VO_2_ measurements for each behavior chamber were divided by mass. A respiratory quotient (RQ) was calculated using the following equation: VCO_2_ output / VO_2_ consumption.

### Statistical Analysis

Prior to running statistical tests, normality was assessed visually from a QQ plot. To assess differences in sex or age on sleep and metabolic rate, a two-way ANOVA was performed (factor 1: time of day; factor 2: sex or age). To assess differences in specific sleep or metabolic traits, a T-test was performed. To assess differences in feeding treatment on sleep and metabolic rate, a one-way ANOVA was performed. To characterize the relationship between the change in CO_2_ output and bout length, we performed linear regression analyses. An F-test was used to determine whether the slope of each regression line was different from zero, while an ANCOVA was used to compare the slopes of different treatments. All *post hoc* analyses were performed using Sidak’s multiple comparisons test. Statistical analyses were performed using GraphPad Prism 9.1 (GraphPad Software). Where represented in each figure, all bars indicate mean values, error bars indicate SEM, and gray circles indicate individual data points.

## RESULTS

To measure the whole-body metabolic rate of individual flies during wake and sleeping states, we developed the SAMM system, a stop-flow indirect calorimetry system that simultaneously uses infrared breams to record activity in freely moving flies (Figure 1). In order to increase the sensitivity of the system over a previously published version (Brown et al., 2019; Botero et al., 2021; Stahl et al., 2017; Stahl et al., 2018; Zandawala et al., 2018), we included an advanced analysis of the raw data consisting of smoothing of the flow rate and CO_2_ data as well as drift correction (Table 1). We also utilized zero-grade air, which contains less than 0.1 ppm total hydrocarbons, to minimize the amount of ambient CO_2_ entering the respirometry system. Overall, these modifications effectively decreased the baseline VCO_2_ measurements from ∼20 μL/hr to ∼3 μL/hr over those previously published (Stahl et al., 2017). We then sought to test the ability of this system to quantify metabolic rate under a variety of contexts.

**Table 1.** List of modifications applied to the analysis of CO_2_ output in the SAMM system.

| **Application** | **Previous System** | **New System** |
| --- | --- | --- |
| Lag Correction | Shifted 8 sec | Shifted 8 sec |
| Smoothing (flow rate) | Rectangular transformation | Savitsky-golay transformation |
| Smoothing (CO<sub>2</sub>) | n/a | Savitsky-golay transformation |
| Drift Correction | Two-point drift correction | Marker baseline correction with the Catmull-Rom Spline option |

The initial version of this system detected changes in CO_2_ output across the circadian cycle, but was unable to resolve sex-specific differences. To test the ability of the updated system to measure changes CO_2_ output associated with these variables, we compared CO_2_ output across the circadian cycle in male and female flies (Figure 2A). In the updated system, female flies sleep significantly more at night than male flies (Figure 2B,C). Further, CO_2_ output was significantly greater during both the day and night in female flies, compared to male flies (Figure 2D-F). Therefore, the updated system is capable of reliably identifying sex differences across the circadian cycle.

**Figure 2.**
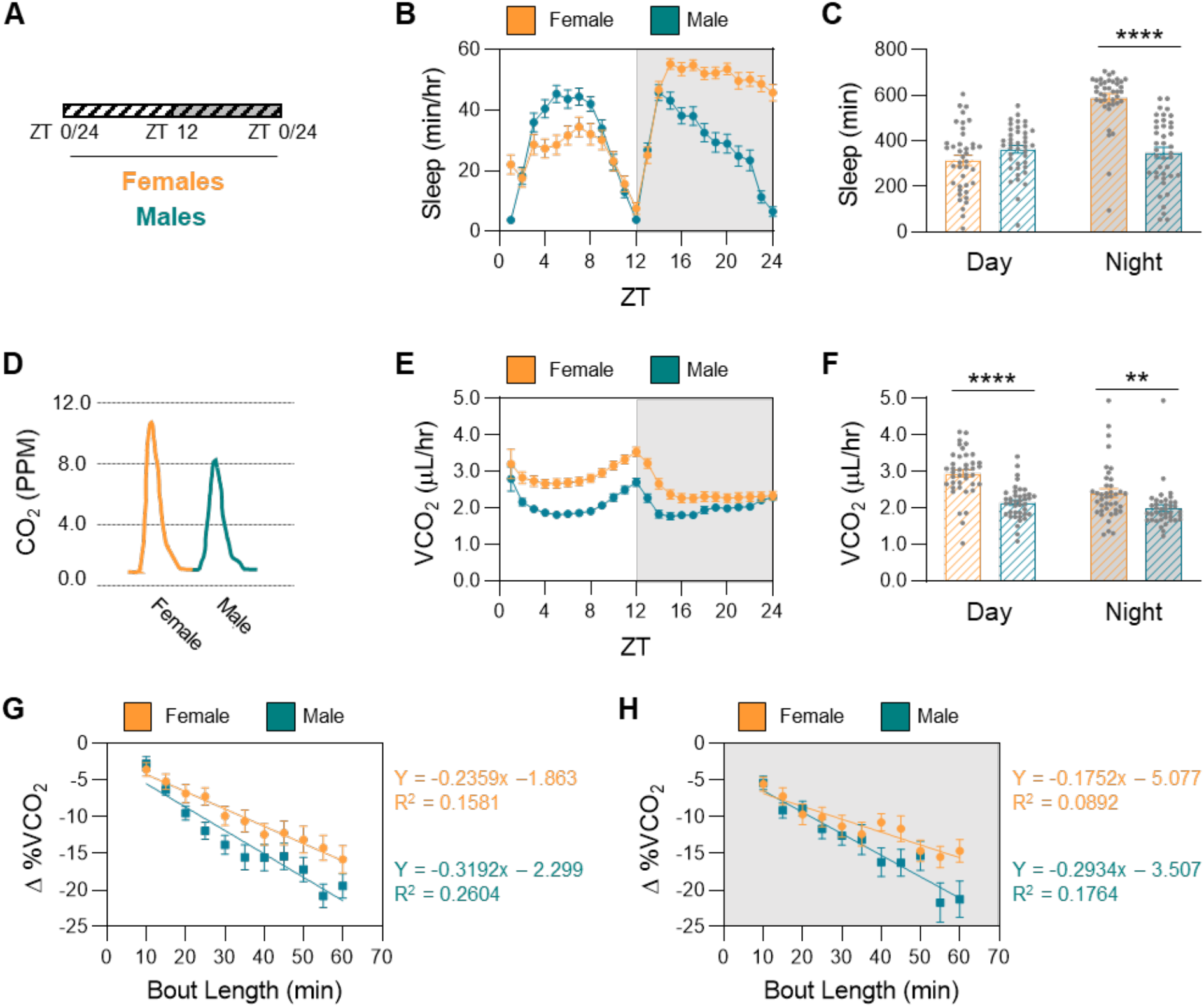
Nighttime sleep and CO_2_ output are reduced in male flies. (**A**) Schematic of experimental design. Sleep and CO_2_ output were assessed in male and female flies for 24 hrs on 5% sucrose. ZT=0 indicates time of lights on, while ZT=12 indicates time of lights off. (**B**) Sleep profile of female and male flies. (**C**) There is a significant effect of sex (two-way ANOVA: F_1,156_ = 21.32, *P*<0.0001, N = 40 per sex) and time of day (two-way ANOVA: F_1,156_ = 39.00, *P*<0.0001, N = 40 per sex) on sleep duration. Females sleep significantly more at night (*P*<0.0001), while no significant differences in sleep duration between day and night was observed in males (*P*=0.8362). Females also sleep significantly more than males during the night (*P*<0.0001), but no significant differences in sleep duration was observed between the sexes during the day (*P*=0.1798). (**D**) Representative trace from an individual female and male, indicating the unadjusted amount of CO_2_ produced over a 5 min time period. (**E**) Profile of CO_2_ output of female and male flies. (**F**) There is a significant effect of sex (two-way ANOVA: F_1,156_ = 38.55, *P*<0.0001, N = 40 per sex) and time of day (two-way ANOVA: F_1,156_ = 12.47, *P*=0.0005, N = 40 per sex) on CO_2_ output. CO_2_ output is significantly higher in females during the day, relative to the night (*P*=0.0003), while no significant differences in CO_2_ output between day and night was observed in males (*P*=0.4760).CO_2_ output is also significantly higher in females both during the day (*P*<0.0001) and night (*P*=0.0065), compared to males. (**G**) Linear regression of daytime CO_2_ output as a function of bout length for both males and females. The slope of each regression is significantly different from zero (females: F_1,399_ = 74.92, *P*<0.0001; males: F_1,390_ = 137.3, *P*<0.0001) as well as significantly different from each other (ANCOVA with bout length as the covariate: F_1,789_ = 4.682, *P*=0.0308). (**H**) Linear regression of nighttime CO_2_ output as a function of bout length for both males and females. The slope of each regression is significantly different from zero (females: F_1,432_ = 43.32, *P*<0.0001; males: F_1,376_ = 80.53, *P*<0.0001) as well as significantly different from each other (ANCOVA with bout length as the covariate: F_1,808_ = 7.925, *P*=0.0050). For profiles and linear regressions, white background indicates daytime, while gray background indicates nighttime. ZT indicates zeitgeber time. Error bars represent +/-standard error from the mean. Grey dots represent measurements of individual flies. ** = *P*<0.01; **** = *P*<0.0001.

To investigate how CO_2_ output changes over the length of a sleep bout, we performed linear regression analyses. We previously reported that, over the course of a 60-min sleep bout, CO_2_ output in females is reduced by ∼15 % and find similar results here (Stahl et al., 2017). We found that during both the day and night, both females and males significantly reduce their CO_2_ output with increasing bout length, as the slope of their respective regression lines were significantly different from zero (Figure 2F,G). A comparison of the two regression lines revealed a significantly greater decrease in CO_2_ output in males (ANCOVA), suggesting that both the magnitude and rate of change in CO_2_ output is greater in males than in females, raising the possibility that there are sex-specific differences in sleep-metabolism interactions.

To determine whether the SAMM system is capable of detecting naturally-occurring differences in sleep quantity and quality, we measured sleep and CO_2_ output in young and aged female flies (Figure 3A). In *Drosophila*, aging results in fragmented sleep that is indicative of diminished sleep quality (Koh et al., 2006; Vienne et al., 2016). In agreement with previous findings using *Drosophila* Activity Monitors (DAMs; Koh et al., 2006; Metaxakis et al., 2014) sleep duration was reduced during the day and night in 40 day-old female flies, compared to 10 day-old female flies (Figure 3B,C). This decrease in sleep duration was accompanied by a significant decrease in bout length and increase in bout number (Figure 3D-E), suggesting an increase in sleep fragmentation in aging flies. Further, aging is also associated with an increase in locomotor activity (White et al., 2010). Towards this end, we found that waking activity is also significantly increased in the aging flies in the SAMM system (Figure 3F). Therefore, many of the previously identified changes in sleep and activity associated with aging are present in the SAMM system, despite differences in arena size and air flow when compared to standard *Drosophila* activity monitors.

**Figure 3.**
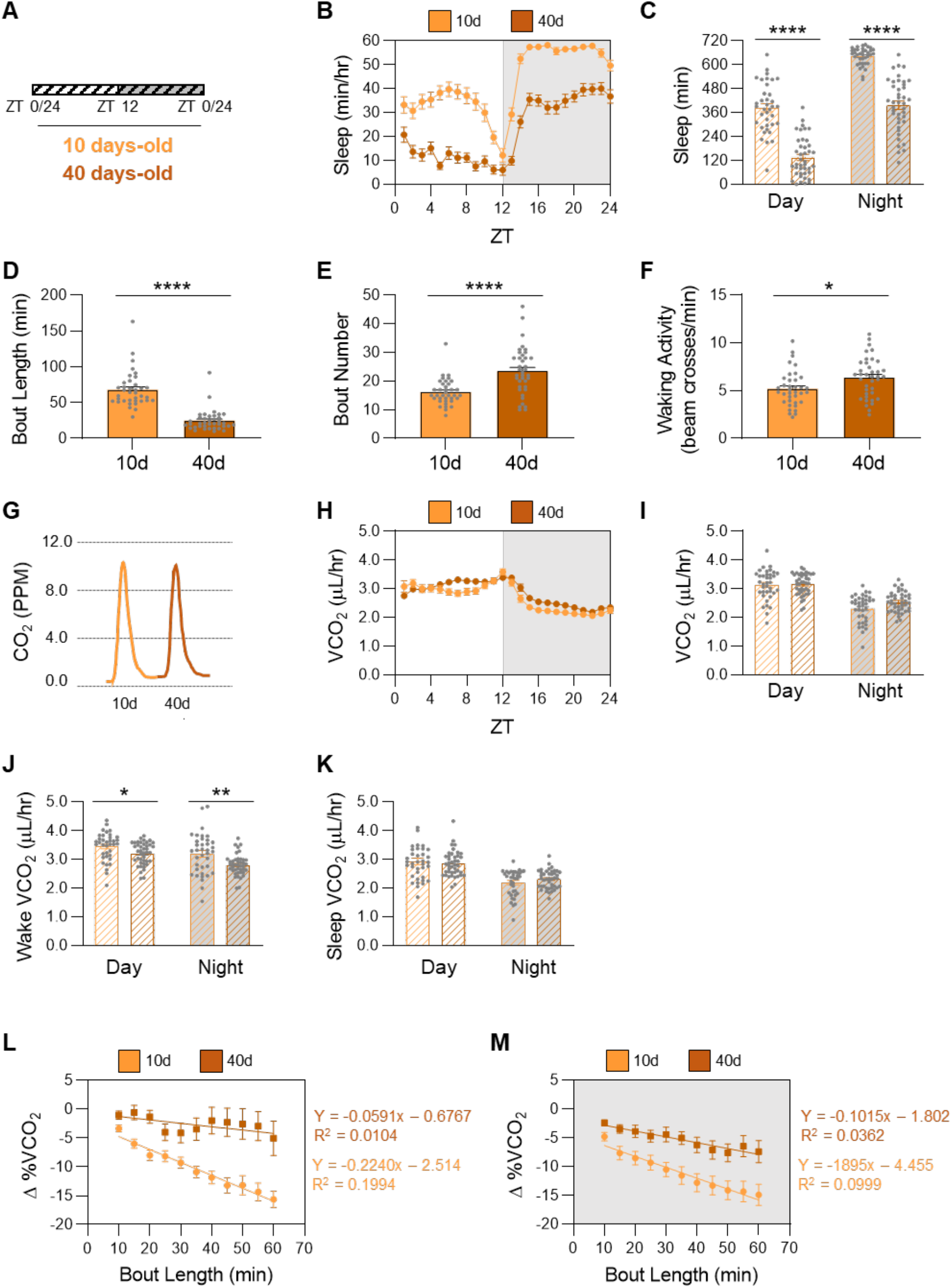
Aging reduces sleep duration and metabolic rate during waking in female flies. (**A**) Schematic of experimental design. Sleep and CO_2_ output were assessed in young, 10 day-old flies and old 40-day old flies for 24 hrs on 5% sucrose. ZT=0 indicates time of lights on, while ZT=12 indicates time of lights off. (**B**) Sleep profile of 10 day-old flies and 40 day-old flies. (**C**) There is a significant effect of age (two-way ANOVA: F_1,152_ = 213.4, *P*<0.0001, N = 37-41) and time of day (two-way ANOVA: F_11520_ = 235.6, *P*<0.0001, N = 37-41) on sleep duration. For both ages, sleep duration is significantly higher during the night, relative to the day (10d: *P*<0.0001; 40d: *P*<0.0001). Sleep duration is significantly reduced in 40 day-old flies both during the day (*P*<0.0001) and night (*P*<0.0001), compared to 10 day-old flies. (**D**) Bout length is significantly reduced in 40 day-old flies, compared to 10 day-old flies (t-test: t_73_=9.318, *P*<0.0001). (**E**) Bout number is significantly increased in 40 day-old flies, compared to 10 day-old flies (t-test: t_73_=04.659, *P*<0.0001). (**F**) Waking activity is significantly increased in 40 day-old flies, compared to 10 day-old flies (t-test: t_73_=2.558, *P*=0.0126). (**G**) Representative CO_2_ trace from an individual 10 day-old and 40 day-old fly. (**H**) Profile of CO_2_ output of young, 10 day-old flies and old, 40 day-old flies. (**I**) There is no effect of age on CO_2_ output (two-way ANOVA: F_1,152_ = 3.155, *P*=0.0777, N = 37-41), but there is an effect of time of day (two-way ANOVA: F_1,152_ = 108.3, *P*<0.0001, N = 37-41) on CO_2_ output. At both ages, CO_2_ output is significantly higher during the day, relative to the night (10d: *P*<0.0001; 40d: *P*<0.0001). (**J**) When awake, there is a significant effect of age (two-way ANOVA: F_1,154_ = 15.90, *P*<0.0001, N = 37-41) and time of day (two-way ANOVA: F_1,154_ = 17.03, *P*<0.0001, N = 37-41) on CO_2_ output. For both ages, CO_2_ output is significantly higher during the day, relative to the night (10d: *P*<0.0457; 40d: *P*<0.0007). CO2 output is also significantly reduced in 40 day-old flies both during the day (*P*<0.0487) and night (*P*<0.0015), compared to 10 day-old flies. (**K**) When sleeping, there is no effect of age (two-way ANOVA: F_1,154_ = 1.794, *P*<0.1827, N = 37-41), but there is a significant effect of time of day (two-way ANOVA: F_1,154_ = 69.18, *P*<0.0001, N = 37-41) on CO_2_ output. For both ages, CO_2_ output is significantly higher during the day, relative to the night (10d: *P*<0.0001; 40d: *P*<0.0001). (**L**) Linear regression of daytime CO_2_ output as a function of bout length for 10 day-old and 40 day-old flies. The slope of 10 day-old flies is significantly different from zero (F_1,420_ = 104.6, *P*<0.0001), while the slope of 40 day-old flies is not (F_1,304_ = 3.191, *P*<0.0751). They are, however, significantly different from each other (ANCOVA: F_1,724_ = 18.55, *P*<0.0001). (**M**) Linear regression of nighttime CO_2_ output as a function of bout length for 10 day-old and 40 day-old flies. The slope of each regression is significantly different from zero (10d: F_1,427_ = 47.37, *P*<0.0001; 40d: F_1,475_ = 17.85, *P*<0.0001) as well as significantly different from each other (ANCOVA: F_1,902_ = 5.841, *P*=0.0159). For profiles and liner regressions, white background indicates daytime, while gray background indicates nighttime. ZT indicates zeitgeber time. Error bars represent +/-standard error from the mean. Grey dots represent measurements of individual flies. * = *P*<0.05; ** = *P*<0.01; *** = *P*<0.001; **** = *P*<0.0001.

In mammals, aging is associated with a reduction in total metabolic rate (Krems et al., 2005; St-Onge & Gallagher, 2010), yet previous work suggests metabolic rate does not change with age in a number of *Drosophila* species tested (Hulbert et al., 2004; Promislow & Haselkorn, 2002). In the SAMM system, total CO_2_ output did not significantly differ between young and old female flies (Figure 3G-I). To specifically examine the effects of sleep on CO_2_ output, we compared the overall CO_2_ output during waking and sleep. We found that CO_2_ output during waking was significantly reduced in aged flies during both the day and night (Figure 3J), while no significant difference in CO_2_ was observed during sleep (Figure 3K). Therefore, although total CO_2_ output doesn’t differ in aged flies, a systematic dissection of CO_2_ output into sleep/waking states revealed a reduction in CO_2_ output during the waking period. This is likely reflective of subtle changes in the circadian regulation of metabolic rate that are obscured in analysis of the daily average metabolic rate.

To directly test whether sleep-metabolism interactions are disrupted in aged flies, we measured CO_2_ output over the length of a sleep bout. We found that during the day, 10 day-old flies significantly reduce their CO_2_ output with increasing bout length, as the slope of the regression line was significantly different from zero (slope = -0.224). However, this is not the case with 40-day old flies, where the slope of the regression line was not significantly different from zero (slope = -0.059; Figure 3L). A comparison of the two regression lines revealed a significantly greater decrease in CO_2_ output in 10 day-old flies, compared to 40 day-old flies (ANCOVA). During the night, both 10- and 40 day-old flies significantly reduce their CO_2_ output with increasing bout length, since the slopes of both regression lines were significantly different from zero (10d: slope = -0.189; 40d: slope = -0.101; Figure 3M). Again, a comparison of the two regression lines revealed a significantly greater decrease in CO_2_ output in 10 day-old flies, compared to 40 day-old flies (ANCOVA). These findings suggest that sleep-dependent changes in CO_2_ output are diminished in aged flies, thereby providing a physiological readout for diminished sleep quality in aged flies.

The initial version of the SAMM was used to investigate CO_2_ output in single flies. Here, we sought to determine whether the system is capable of identifying context-specific changes in O_2_ consumption. The previous version of the SAMM system was insufficiently sensitive to detect O_2_ levels in single flies, so experiments were performed in groups of 25 flies. To measure O_2_ consumption, we included analysis using an O_2_ analyzer (Figure 1). First, CO_2_ levels were measured as described above. The reference and animal air were next passed through scrubbing columns to remove both humidity and CO_2_, since these factors influence O_2_ levels, and then were sent to the O_2_ analyzer. Similar to analyses of single flies, we found that CO_2_ output was significantly increased during the day in both sexes (Figure 4A,B), suggesting the process does not disrupt the relative readings of CO_2_ levels. Using this setup, we found that O_2_ consumption is also significantly higher during the day in both sexes (Figure 4C,D), suggesting the SAMM system has sufficient sensitivity to detect changes in O_2_ levels in groups of flies. It is possible that this increase in CO_2_ output and O_2_ consumption in females may be a consequence of inherent differences in body size between the sexes, with females generally being larger than males. To address this possibility, we adjusted the CO_2_ and O_2_ measurements from each behavior chamber for weight. We found that CO_2_ output was significantly increased in males during both the day and night (Supplementary Figure 1A,B). Similarly, we found that O_2_ consumption is also significantly higher in males during both the day and night (Supplementary Figure 1C,D), Overall, these results suggest that males have higher basal metabolic rates than females.

**Figure 4.**
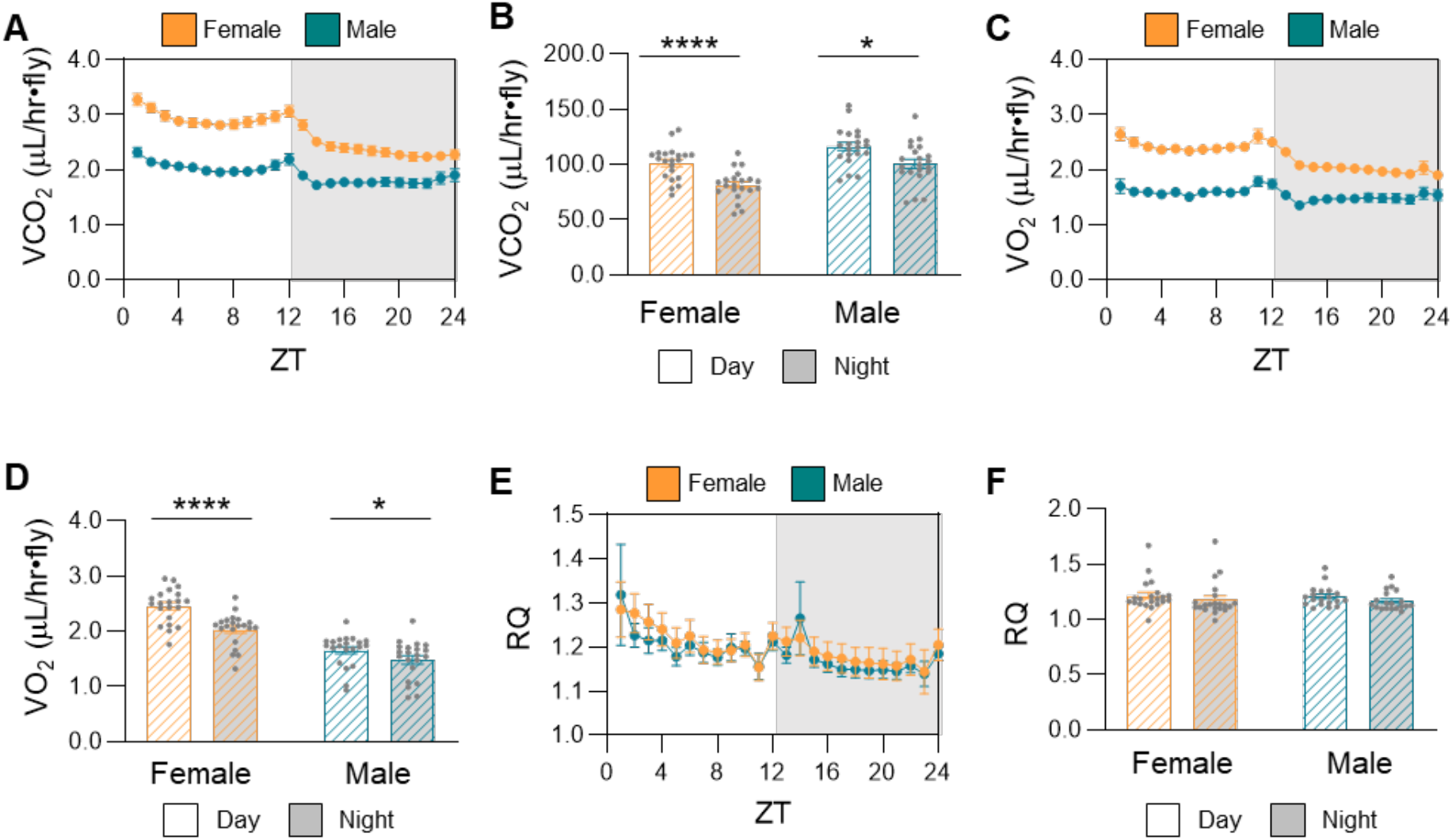
The Sleep, Activity, and Metabolic Monitor (SAMM) system can be used to measure CO_2_ output and O_2_ consumption in groups of flies. Groups of 25 flies were fed 5% sucrose, during which both CO_2_ and O_2_ were measured, as described in Figure 1, over the course of 24 hours. (**A**) Profile of CO_2_ output in female and male flies. (**B**) There is a significant effect of sex (two-way ANOVA: F_1,80_ = 85.25, *P*<0.0001, N = 21 per sex) and time of day (two-way ANOVA: F_1,80_ = 29.24, *P*<0.0001, N = 21 per sex) on CO_2_ output. For both sexes, CO_2_ output is significantly higher during the day, relative to the night (females: *P*<0.0001; males: *P*=0.0301). CO_2_ output is also significantly higher in females both during the day (*P*<0.0001) and night (*P*<0.0001), compared to males. (**C**) Profile of O_2_ consumption in female and male flies. (**D**) There is a significant effect of sex (two-way ANOVA: F_1,80_ = 95.36, *P*<0.0001, N = 21 per sex) and time of day (two-way ANOVA: F_1,80_ = 18.72, *P*<0.0001, N = 21 per sex) on O_2_ consumption. For both sexes, O_2_ consumption is significantly higher during the day, relative to the night (females: *P*<0.0001; males: *P*=0.0125). O_2_ consumption is also significantly higher in females both during the day (*P*<0.0001) and night (*P*<0.0001), compared to males. (**E**) Profile of the respiratory quotient (CO_2_ output / O_2_ consumption) in female and male flies. (**F**) There is no significant difference in the respiratory quotient between male and females (two-way ANOVA: F_1,80_ = 0.1353, *P*=0.7140, N = 21 per sex). For profiles, white background indicates daytime, while gray background indicates nighttime. ZT indicates zeitgeber time. Error bars represent +/-standard error from the mean. Grey dots represent measurements of individual flies. **** = *P*<0.0001.

Given that both the total CO_2_ output and total O_2_ consumption are known, the metabolic fuel being utilized can be estimated based on the respiratory quotient (RQ; Ferrannini, 1988), which is the ratio of VCO_2_ to VO_2_. In animals, when RQ = 1, carbohydrates are the primary fuel for metabolic processes; when RQ<1, fat and protein are utilized (Kleiber, 1962); and when RQ>1, fats are synthesized from carbohydrates to be used as fuel (Prinzinger et al., 1992; Talal et al., 2020; Voigt & Winter, 1999). We found that both males and females, despite differences in metabolic rate, have similar RQ on the all-sucrose diet used in our assay system, with an RQ of ∼1.2 for both sexes during the day and ∼1.17 for both sexes during the night (Figure 4E,F), suggesting that dietary sugar provided to the flies is being converted to fat. Although it appears that the RQ is elevated during the first ∼4 hrs after lights on (ZT 0-4), when binned into 4 hour intervals, there are no significant differences between bins (data not shown). Therefore, for both males and females, the RQ remains constant across the circadian cycle, despite overall changes in total CO_2_ release and O_2_ consumption. Overall, the SAMM system can reliably detect CO_2_ output and O_2_ consumption in groups of flies, providing a system to examine the genetic basis of metabolic rate and energy utilization.

## DISCUSSION

Measurements of whole-body metabolic rate are widely used to study physiological changes associated with environmental and life-history traits across phyla (Chabot et al., 2016; Clarke et al., 2010; Gardner et al., 2020; Jetz et al., 2008; Nagy, 2005; Norin & Clark, 2016), including *Drosophila* and other insects (Contreras & Bradley, 2010; Reinhold, 1999). Here, we describe updates to the SAMM system, which allows us to (1) simultaneously measure sleep-wake activity and metabolic rate in single flies and (2) estimate nutrient utilization from respiratory quotients in groups of flies. Our system provides precise measurements of metabolic rate and activity across the circadian cycle. We are able to record from five individual flies from a single CO_2_ analyzer through indirect calorimetry, making this a moderate throughput system. Using this updated system, we have validated our previous findings and extend analysis to characterize sex- and age-specific differences in metabolic rate.

Multiple systems have been applied to measure whole body metabolic rate in flies. For example, there are technically simple systems that measure of CO_2_ production based on indirect fluid displacement (Yatsenko et al., 2014; Kucherenko et al., 2011). This approach is cost effective and high-throughput. Another paradigm to measure metabolic rate includes feeding radiolabeled substrates to flies and then measuring the exhaled radiolabeled CO_2_ that is produced (Bland, 2016; Francis et al., 2019). However, these approaches are unlikely to be sufficiently sensitive to measure CO_2_ production in single flies and have low temporal resolution. Indirect calorimetry has been widely used in *Drosophila* to measure CO_2_ production (e.g. Botero et al., 2021; Jensen et al., 2014; Nagarajan-Radha et al., 2020; Stahl et al., 2018). In these studies, CO_2_ release from individuals or groups of flies can be directly measured using a CO_2_ analyzer, providing a close to real time measurement of CO_2_ production. This approach has been used to test the effects of many different environmental factors including aging (Jensen et al., 2014; Khazaeli et al., 2005; Promislow & Haselkorn, 2002), as well as genetic mutations and backgrounds (Botero et al., 2021; Nagarajan-Radha et al., 2020; Stahl et al., 2017; Stahl et al., 2018; Zandawala et al., 2018). The SAMM system uses a stop-flow configuration that increases sensitivity, allowing for detection of CO_2_ in single flies, while simultaneously measuring activity. Direct calorimetry has also been utilized in *Drosophila*, in which metabolic activity is measured by heat output. Towards this end, a system was recently developed that simultaneously measures heat production and activity, providing many of the functional advantages of the SAMM system (Fiorino et al., 2018). Therefore, multiple systems are available to measure sleep-wake activity and metabolic rate in flies.

The ability to simultaneously record CO_2_ and O_2_ levels in group-housed flies provide the opportunity to systematically investigate the mechanisms underlying metabolism of different energy stores. Our analysis found no difference in RQ between female and male flies when fed a sucrose diet. A number of other studies have examined the RQ in insects, but surprisingly little is known about how diet, genes, and life-history traits influence macronutrient metabolism. Using a similar respirometry system, a recent study has shown that the RQ approaches 1 in female flies fed standard media, and decreases (0.8) during starvation (Wat et al., 2020). In locusts, the RQ increases in diets with a high carbohydrate:protein ratio to 1.2 (Talal et al., 2020). In this study, as in previous studies using the SAMM, flies were raised on standard diet, then fed sugar-alone for testing (Brown et al., 2019; Stahl et al., 2017; Zandawala et al., 2018). Because sugar-only diets have been shown to influence sleep-depth based on measurements of arousal threshold (Brown et al., 2020), it is possible that systematic analysis of diet throughout development, or acutely during metabolic testing will reveal differences in respiratory quotient.

The ability to precisely measure sleep-dependent changes in metabolic rate may provide a tractable way to examine physiological correlates of sleep intensity that have previously been measured through arousal threshold and neural activity (Shafer & Keene, 2021). This is critical as the vast majority of sleep studies in *Drosophila* have used behavioral analysis. These studies have clearly identified sex-specific differences in sleep, but it is less clear how they impact sleep intensity (Garbe et al., 2016; Harbison & Sehgal, 2008; Isaac et al., 2010). Similarly, sleep-dependent changes in aging have been identified in flies including fragmented bout length in aged flies (Koh et al., 2006; Vienne et al., 2016). The SAMM system identifies sex- and age-specific differences in metabolic rate. For example, a decrease in metabolic rate of males during longer sleep bouts suggests they sleep more deeply, and could explain reduced nighttime sleep in male flies compared to females. Conversely in aged flies, the sleep-dependent decrease in metabolic rate is reduced, suggesting aging disrupts sleep depth and/or sleep-metabolism interactions. Extending genetic analysis of age- and sex-specific differences in sleep that have previously been studied using behavioral readouts to physiological analysis will likely uncover complexities in the impact of sex and life history on sleep regulation that have previously been overlooked.

## CONCLUSIONS

Here, we provide a detailed description of the updated SAMM system to simultaneously measure sleep-wake activity and metabolic rate in single flies. This system is largely comprised of commercially available components and therefore could be readily replicated in other laboratories. We provide proof-of-principle experiments for the use of this system to identify age- and sex-specific changes in metabolic rate. This system, combined with the ability to manipulate gene and neural function, and a rapidly increasing understanding of the biological mechanisms underlying sleep and metabolic regulation, has potential to advance our understanding of sleep-metabolism interactions.

## CREDIT AUTHORSHIP CONTRIBUTION STATEMENT

Conceptualization: Elizabeth B. Brown, Alex C. Keene, Jaco Klok

Data curation: Elizabeth B. Brown, Jaco Klok

Formal analysis: Elizabeth B. Brown

Funding acquisition: Alex C. Keene, Elizabeth B. Brown

Investigation: Elizabeth B. Brown

Methodology: Elizabeth B. Brown, Jaco Klok

Project administration: Alex C. Keene

Resources: Jaco Klok, Alex C. Keene

Software: Jaco Klok

Supervision: Alex C. Keene

Validation: Elizabeth B. Brown

Visualization: Elizabeth B. Brown

Writing – original draft: Elizabeth B. Brown, Alex C. Keene

Writing – review & editing: Elizabeth B. Brown, Alex C. Keene, Jaco Klok

## ACKNOWLEDGEMENTS

We are thankful to members of the Keene laboratory for helpful discussions and technical support. This work was supported by the National Institutes of Health [grant numbers R21NS124198 and R01DC017390 to A.C.K, and K99AG071833 to E.B.B].

## FIGURE LEGENDS

**Supplementary Figure 1.** Weight-adjusted measurements of CO_2_ output and O_2_ consumption in groups of flies. CO_2_ output and O_2_ consumption measurements were obtained from Figure 4. (**A**) Profile of CO_2_ output in female and male flies, adjusted by weight. (**B**) When adjusted for weight, there is a significant effect of sex (two-way ANOVA: F_1,80_ = 23.14, *P*<0.0001, N = 21 per sex) and time of day (two-way ANOVA: F_1,80_ = 25.08, *P*<0.0001, N = 21 per sex) on CO_2_ output. For both sexes, CO_2_ output is significantly higher during the day, relative to the night (females: *P*<0.0003; males: *P*=0.0049). CO_2_ output is also significantly higher in males both during the day (*P*<0.0074) and night (*P*<0.0005), compared to males. (**C**) Profile of O_2_ consumption in female and male flies, adjusted by weight. (**D**) When adjusted for weight, there is a significant effect of sex (two-way ANOVA: F_1,80_ = 13.62, *P*<0.0004, N = 21 per sex) and time of day (two-way ANOVA: F_1,80_ = 7.460, *P*<0.0078, N = 21 per sex) on O_2_ consumption. For both sexes, O_2_ consumption is significantly higher during the day, relative to the night (females: *P*<0.0103; males: *P*=0.0429). O_2_ consumption is also significantly higher in males both during the day (*P*<0.0137) and night (*P*<0.0023), compared to females. For profiles, white background indicates daytime, while gray background indicates nighttime. ZT indicates zeitgeber time. Error bars represent +/-standard error from the mean. Grey dots represent measurements of individual flies. **** = *P*<0.0001.

