## Supplementary figures and images for "Measuring metabolic rate in single flies during sleep and waking states"

### Supplemental Figure 1

Supplementary Figure 1

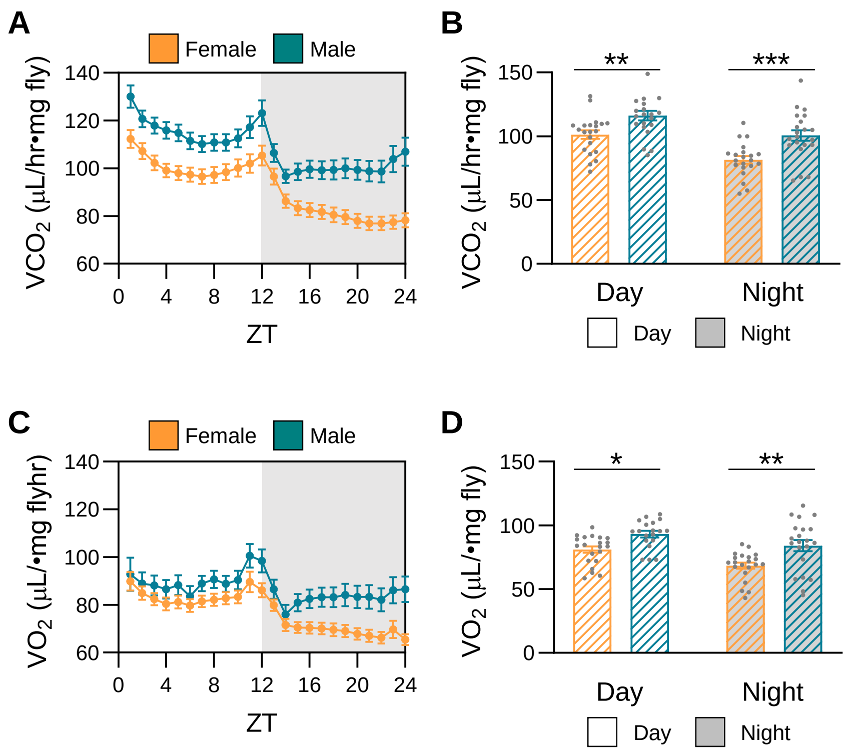
